# RecQ1 helicase unwinds G-Quadruplexes at oriLyt to facilitate KSHV Lytic DNA Replication

**DOI:** 10.1101/2022.12.26.521964

**Authors:** Prerna Dabral, Timsy Uppal, Subhash C. Verma

## Abstract

KSHV utilizes distinct origins of DNA replication (origin of replications) during the latent and lytic phases of its life cycle. Lytic DNA replication requires the participation of cellular and viral proteins to initiate replication at a specific region in the KSHV genome, oriLyt. These regions contain multiple secondary structures and binding sites for the viral and cellular proteins. We discovered the presence of G-quadruplex (G4s) sites in the oriLyt region. G-quadruplexes are secondary structures in nucleic acid sequences that are considered regulators of multiple biological processes, such as transcription, translation, and replication initiation. Importantly, herpesviruses have a high density of PQS (putative quadruplex formation sites) in their genome, including in the regulatory regions, which control replication and transcription. The binding of RecQ1 to the oriLyt region of KSHV and its ability to unwind the G4 structures led us to speculate that G-quadruplex sites play an important role in lytic DNA replication. In this study, we confirmed the formation of stable G-quadruplexes through biochemical and biophysical assays. We further demonstrated that RecQ1 associates with the G4 sites of the oriLyt. The functional significance of RecQ1-mediated unwinding of G4 sites was confirmed by the inhibition and depletion of RecQ1 activity and protein levels through a pharmacologic inhibitor and short-hairpin, respectively. Furthermore, the detection of replication initiation through single-molecule analysis of the replicated DNA (SMARD) approach demonstrated that G4 stabilization leads to a subdued initiation of replication at the oriLyt. This confirmed the role of the G-quadruplex in regulating viral DNA replication, which can be used for controlling virus growth.

**Significance:** Biological processes originating from the DNA and RNA can be regulated by the secondary structures present in the stretch of nucleic acids, and the G-quadruplexes are shown to regulate transcription, translation, and replication. In this study, we identified the presence of multiple G-quadruplex sites in the region (oriLyt) of KSHV DNA, which is essential for the replication of DNA during the lytic cycle. We demonstrated the roles of these G-quadruplexes through multiple biochemical and biophysical assays in controlling replication and efficient virus production. We demonstrated that KSHV achieves this by recruiting RecQ1 (helicase) at these G-quadruplex sites for efficient viral DNA replication. Analysis of the replicated DNA through nucleoside labeling and immunostaining showed a reduced initiation of DNA replication in cells with a pharmacologic stabilizer of G-quadruplexes. Overall, this study confirmed the role of the G-quadruplex in regulating viral DNA replication, which can be exploited for controlling viral DNA replication.

## Introduction

Kaposi Sarcoma associated Herpesvirus (KSV) or HHV-8, a gammaherpesvirus, is the etiological agent of Kaposi’s Sarcoma, Primary effusion lymphoma, Multicentric Castleman’s disease, and KSHV inflammatory syndrome (KICS) (1–4). Like other viruses of the Herpesviridae family, the life cycle of KSHV is biphasic, characterized by a default latent phase and a short and transient lytic phase. The latent phase of the viral life cycle is characterized by restricted gene expression and latent DNA replication. Lytic DNA replication occurs once per cycle, facilitated by the latent associated nuclear antigen (LANA) after recruiting the cellular replication machinery to the origin of latent DNA replication (5). The lytic phase of infection involves an orchestrated expression of immediate early (IE), early (E), and late (L) genes with the replication of the viral genome to produce infectious virions following packaging into capsids and teguments (6). Lytic replication initiates at the origin of lytic DNA replication (oriLyt), during which the virus uses viral and cellular proteins to replicate its genome, presumably through a rolling circle mechanism. Two copies of oriLyt in the KSHV genome, oriLyt-L and oriLyt-R, are present between the open reading frames (ORFs) K4.2 and K5, and between K12 and ORF71, respectively (7, 8). The oriLyt regions are 1.7Kb in length, consisting of a highly similar 1.1Kb long core sequence along with 600bp GC repeats, which are present in the form of 20bp and 30bp tandem repeats. The core sequence contains C/EBP motifs, which are required for lytic DNA replication through its binding to the K8 protein at the AT palindromic sequence. In addition, the core sequence contains an RTA-dependent promoter, which includes RTA binding site, RRE (RTA response element), and TATA box consensus sequence (9, 10).

Proteins that are essential for lytic DNA replication are conserved among herpesviruses forming the core replication machinery, consisting of ORF9 (DNA polymerase), ORF6 (single-stranded binding protein), ORF59 (processivity factor), ORF44 (helicase), PRF56 (primase), and ORF 40/41 (primase-associated factor) (11–13). ORF50 or RTA and K8 are essential for lytic DNA replication and have been shown to form the pre-replication complex with the core proteins following recruitment to the oriLyt region (9, 14).

In addition to the viral proteins, many cellular proteins were identified to bind to the oriLyt region, including (Topo) I and II, MSH2/6, RecQL, DNA-PK, poly (ADP-ribose) polymerase 1 (PARP-1), Ku autoantigens and SAF-A (scaffold attachment factor A). These proteins are involved in several cellular processes, such as replication, recombination, and repair, and are shown to be present on the viral replication foci during lytic replication.

G-quadruplexes or G4 structures are thermodynamically stable non-canonical structures formed in DNA/RNA sequences carrying four runs of at least three guanines separated by a few bases (15). The building block of a G-quadruplexes is a guanine tetrad, formed by hoogsteen hydrogen bonding between four guanines giving rise to a planar structure that stacks on top of each other through π-π interactions. The presence of monovalent cations such as K+ ions and Na+ ions reduces repulsion between negatively charged oxygen atoms .thereby, promoting the formation of G-quadruplexes (16). These structures have been identified in eukaryotes, prokaryotes, and viruses and are implicated in regulating various biological processes, including replication, transcription, and translation (17). The presence of G-quadruplexes has been reported in several human viruses, including Human Immunodeficiency Virus (HIV), Hepatitis B virus, Hepatitis C virus, Ebola Virus, Zika virus, Epstein Barr Virus (EBV), and KSHV (18, 19).

Recently, G-quadruplexes became important regulators of replication initiation at the origin sites (20). These structures have been identified in approximately 90% of the replication origins in humans and mice, and about 70% of the origins/replication initiation sites in Drosophila are reported to be a few hundred base pairs downstream of G4 sites (21). G-quadruplex regions have been identified as the potential site of replication initiation, and their roles are demonstrated in the early replication (22, 23). Moreover, the association of G-quadruplexes with proteins involved in their unwinding or resolving negatively supercoiled duplex DNA has led to the hypothesis that the unfolding of these secondary structures located in origin facilitates the recruitment of many replication factors such as ORCs, which have affinities for these G4 sites (22, 24).

Moreover, the presence of G-quadruplexes in the genome could be detrimental, as they serve as roadblocks to the movement of cellular enzymes involved in replication or transcription. Therefore, cells employ ways to resolve these structures for the precise functioning of biological processes. Helicases are the cellular motor proteins that separate two annealed strands of nucleic acids using energy derived from the nucleoside triphosphate hydrolysis (25–28). Their role in unwinding G4s was initially reported in the late 1990s, with SV-40 helicase, Bloom syndrome helicase, *Saccharomyces cerevisiae* proteins, Sgs1p helicase, and SEP1 protein reported to resolve the G4 structures (29–32). RecQ belongs to a conserved family of DNA helicases called guardians of the genome due to their roles in DNA replication, recombination, and repair (33, 34). The human RecQ helicases include RECQ1, WRN, BLM, RECQ4, and RECQ5 and Bloom, Werner, and Rothmund-Thompson syndrome, which is associated with mutations in BLM, WRN, and RecQ4, respectively (35). BLM helicase encoded by the BLM gene has been shown to unwind the G4 structures, and its G4 unwinding activity has been implicated in promoting telomere replication and genome integrity (36–39). Another member of the RecQ family, WRN helicase, has been shown to reduce genome instability due to its ability to resolve (CGG) repeats associated with fragile X syndrome in addition to promoting efficient replication at the telomeric sequences (40–42). Recent studies showed that RecQL4, a member of the RecQ family important for replication initiation and fork progression, binds to the G4 structures (43, 44).

RecQ1, another member of the RecQ family, is the most abundant RecQ protein in the cells and gets upregulated in EBV-transformed B cells (45, 46). RecQ1 has been shown to repress chromosome instability, reduce DNA damage and promote initiation at the paused replication forks (47–49). In addition to being linked to replication initiation at the origin, RECQ1 binds to the oriLyt region of KSHV and EBV (50, 51). We previously reported G-quadruplexes’ formation throughout the genome and its impact on the latent DNA replication (52). In this study, we demonstrate the formation of these secondary structures in the oriLyt region through biophysical and biochemical assays.

Moreover, we determined the binding of RecQ1 to the G4 forming region of the oriLyt and substantiated the indispensable role of RecQ1 in KSHV lytic replication through inhibition of RecQ1 helicase activity and depletion of RecQ1. Additionally, we evaluated the relevance of G-quadruplex formation in lytic replication through their disruption by site-direction mutagenesis of the G4 site. We also used ligands to stabilize the formation of these structures to determine their effect on the initiation of lytic replication on individual DNA molecules.

## MATERIALS AND METHODS

### Cell lines, plasmids, and antibodies

The KSHV-positive BCBL-1/BC3 cell lines were grown in RPMI 1640 supplemented with 10% Fetal Bovine Serum (FBS), 2mM glutamine, 5U/ml penicillin, and 5ug/ml streptomycin. BJAB-L-YFP cell line was grown in RPMI 1640 supplemented with 10% FBS, 2 mM glutamine, 5U/ml penicillin and 5ug/ml streptomycin, and 0.5μg/ml puromycin. iSLKTet-RTA-Bac16-wild-type (wt) and iSLKTet-RTA-Bac16-RTA-STOP cells were maintained in DMEM supplemented with 10% tet-Free FBS with additional 600μg/mL hygromycin B, 400μg/mL G418, and 1μg/mL puromycin. iSLK.219 cells with recombinant KSHV BACs were induced by doxycycline. The following commercial antibodies were used for this study: rabbit anti-RecQ1 (Bethyl Laboratories, Inc.), mouse anti-GAPDH (US Biological), and rabbit anti-RTA (custom synthesized at GenScript, Inc.). Site-directed mutagenesis was performed in wt 8088sc (8088-wt) plasmid carrying the KSHV oriLyt-L sequence using QuikChange Lightning Site-Directed mutagenesis kit, according to the manufacturer’s protocol. The integrity of the resulting mutated plasmid, i.e., 8088mut, was confirmed by sequencing at the Nevada Genomics Center, University of Nevada, Reno, NV.

### CD spectroscopy

CD spectroscopy was performed on wt-oriLyt and scrambled oligos using Aviv Biomedical spectrometer, where circularly polarized UV light was used to record CD spectra in a series of progressive scans from 320nm to 200nm in a quartz cuvette of 1mm path length at 25^0^C. The wt- and scrambled oriLyt DNA oligonucleotides were used at a final concentration of 5μM in sodium cacodylate buffer (10mM, pH 7.4) with 100mM KCl. G-quadruplexes were allowed to form in the oligos by initial denaturation at 95°C for 5 minutes, followed by slow cooling at room temperature.

### Electrophoretic mobility shift assay

Electrophoretic mobility shift assay (EMSA) was performed as described before (52). Briefly, wt-oriLyt and scrambled oriLyt DNA oligonucleotides were labeled with radioactive ^32^P using Terminal deoxynucleotide transferase, TdT (New England Biolabs), and resuspended at a final concentration of 2μM in 10mM sodium cacodylate buffer with 100mM KCl. Following denaturation and slow cooling, the oligos were resolved on a 15% native polyacrylamide gel in 1X TBE buffer with 100mM KCl, gels were dried using Gelair gel dryer (Bio-Rad Inc.), and autoradiography was performed using a phosphorimager (GE Healthcare Life Sciences). For G4 disruption, antisense oligos were incubated with the labeled oligos before denaturation.

### *In vitro* DNA pull-down assay

The wt-oriLyt oligo (5’-CACGGGGTTGTTCGGTGGCGGGGGGGGGGCTAG-3’) and scrambled oligo (5’-ACTATATTGTTCAATAACTATATTATAAATACT-3’) were biotinylated at the 3’end Terminal deoxynucleotide transferase, TdT (New England Biolabs). The cellular lysate was prepared from approximately 20 million KSHV-positive BCBL-1 cells, which were lytically induced for 24 hours using sodium butyrate (NaB) and 12-o-tetradecanoylphorbol-13-acetate (TPA), harvested and lysed in 1% NP-40 lysis buffer with protease inhibitors. The samples were sonicated and centrifuged at 4°C to remove cellular debris. The purified biotinylated DNA oligos were incubated with the lysate for three hours at 4°C, and Pierce Streptavidin agarose beads (Thermo Fisher Scientific) were added to the samples for two hours to pull down the proteins bound to the biotinylated DNA. The beads were washed thrice with 1% NP-40 lysis buffer, following which they were loaded onto a 9% SDS-PAGE, transferred to a nitrocellulose membrane, and probed with RecQ1 antibody.

### Chromatin immunoprecipitation assay (ChIP)

Chromatin immunoprecipitation was performed using iDeal ChIP qPCR kit (Diagenode, Inc.) according to the manufacturer’s instructions. Briefly, KSHV-positive cells were induced for lytic reactivation for 24 hours, following which approximately 8 million cells were fixed with 1% formaldehyde for 10 min at room temperature, and cross-linking was stopped by the addition of glycine at a final concentration of 125 mM for 5 min. The cells were washed two times with ice-cold PBS, and nuclei were extracted following cell lysis using kit-supplied buffers. Cells were finally lysed in chromatin shearing buffer supplemented with protease inhibitors for 10 min on ice, and chromatin was sonicated using a Bioruptor (Diagenode) to an average length of 200 to 300 bp. The lysates were centrifuged at 16,000g for 10 minutes to remove the cell debris, and the supernatant was incubated with specific antibodies and magnetic beads overnight at 4 °C. The following day, the beads were collected and subsequently washed at 4 °C for 5 minutes each with wash buffers iW1, iW2, iW3, and iW4. The beads-bound DNA was subjected to Proteinase K digestion, and the DNA was finally eluted from the beads by incubation at 100 °C for 15 minutes. Purified ChIP DNA samples and the inputs were subjected to amplification with specific primers using ABI StepOne plus a real-time PCR machine (Applied Biosystems).

### Transfection of oriLyt plasmids and transient replication assay

Approximately 20 million BCBL-1 cells were transfected with 30 μg of oriLyt plasmids (either 8088wt or 8088mut) using the Neon transfection system (Thermo Fisher Scientific). The cells were allowed to recover for 24 hours and induced lytic reactivation (24 hours for Chromatin Immunoprecipitation assay and 48 hours for replication assay). For replication assay, cells were washed with phosphate-buffered saline followed by extraction of DNA using a modified Hirt’s lysis method, described earlier (53, 54). A fraction of DNA was linearized with EcoRI and the remainder with DpnI and EcoRI to remove the non-replicated DNA. Digested DNA was resolved and transferred to a Nylon membrane, followed by hybridization with ^32^P-labeled 8088sc probes. The auto-radiographic signals were detected using a PhosphorImager, according to the manufacturer’s instructions (Molecular Dynamics, Inc.).

### Single Molecule Analysis of the Replicated DNA (SMARD)

Single Molecule Analysis of Replicated DNA (SMARD) was used to analyze the effect of G-quadruplex formation on the initiation of lytic DNA replication as described previously (55) (56). BCBL-1 cells were treated with G4 stabilizing compound PhenDC-3 and TMPyP4 at a concentration of 10uM and induced lytically for 24 hours. The cells were sequentially labeled with 30 μM 5-iodo-2′-deoxyuridine (IdU) (Sigma-Aldrich) at 37°C for 4 h, washed with PBS, and labeled with second nucleotide analog, 5′-chloro-2′-deoxyuridine (CldU) at 30 µM (Sigma-Aldrich) for 4 h. The cells were finally washed and resuspended in PBS (1 × 10^6^ cells per ml) and molten 1% InCert agarose (Lonza Rockland, Inc., Rockland, ME, USA) in PBS (1:1 vol/vol). Agarose gel plugs embedded with labeled cells were made by solidification in a cold plastic mold on ice for 30 min. The plugs were incubated in lysis buffer (1% *n*-lauroylsarcosine, 0.5 M EDTA, and 20 mg/ml proteinase K) for at least 72 h at 50°C, following which the proteinase K was removed by washing the plugs with 1X TE and phenylmethanesulfonyl fluoride (PMSF) (Sigma-Aldrich). Next, the KSHV genome was linearized using *PmeI* (New England Biolabs Inc.), where the plugs were first washed with pre-digestion buffer (10 mM MgCl_2_ and 10 mM Tris-HCl (pH 8.0) before digesting with 70 units of *PmeI* at 37°C overnight. The *PmeI* digested gel plugs were rinsed twice with TE, casted onto a 0.7% SeaPlaque GTG agarose gel (Lonza Rockland, Inc.), and resolved by PFGE for 36 hours. KSHV genome was detected using Southern blotting and hybridization with a ^32^P-labeled KSHV TR-specific probe. Band specific for KSHV genome was excised from the gel, and DNA was extracted using GELase treatment (Epicentre Biotechnologies, Madison, Wisconsin, 1 unit per 50 μl of agarose suspension) that digests the agarose releasing the DNA into suspension. The DNA was then stretched on 3-aminopropyltriethoxysilane (Sigma-Aldrich) coated glass slides and denatured in alkaline denaturing buffer (0.1N NaOH/70% Ethanol, and 0.1% ß-mercaptoethanol) and fixed with 0.5% glutaraldehyde. The DNA was hybridized overnight with biotinylated probes, and the next day, slides were rinsed in 2x SSC (1x SSC) 1% SDS, washed in 40% formamide solution containing 2x SSC at 45°C for 5 min, and rinsed in 2x SSC-0.1% IGEPAL CA-630. The slides were washed 4 times with 4x SSC-0.1% IGEPAL CA-630, followed by blocking with 1% BSA for 20 min. Next, the slides were treated with NeutrAvidin Alexa Fluor 350 (Life Technologies Inc.) and biotinylated anti-avidin antibodies (Vector Laboratories, Inc.) for 20 min each, after washing with PBS containing 0.03% IGEPAL CA-630 in between. Slides were further treated with NeutrAvidin Alexa Fluor 350 for 20 min and washed as mentioned above. Following this, the slides were incubated with mouse anti-IdU monoclonal antibody, mouse anti-CldU monoclonal antibody, and biotinylated anti-avidin D for 1 h. After washing, the slides were incubated with NeutrAvidin Alexa Fluor 350 and fluorescent secondary antibodies against mouse and rabbit, i.e., Alexa Fluor 488 and Alexa Fluor 594 (Invitrogen Molecular Probes), for one h. The slides were washed again, and coverslips were mounted with ProLong gold anti-fade reagent (Life Technologies Inc.), followed by fluorescence microscopy.

### RecQ1 knockdown using lentiviral vectors

The pTRIPZ lentiviral vector (Dharmacon, GE Life Sciences) containing shRNA for RecQ1 was co-transfected with lentivirus packaging vectors, pCMV-dR8.2 and pCMV-VSVG (Addgene, Inc.) into Lenti-X 293T (Takara Bio) cells using polyethylenimine (PEI) (Polysciences, Inc.) to generate the lentiviral particles. Lentivirus containing supernatants from the transfected cells were collected for 5 days, followed by concentration of the virus by ultracentrifugation (25,000 rpm, 1.5 hr, 4 °C). The concentrated virus was used for transducing the target cells, BCBL-1, in the presence of 5μg/ml polybrene, followed by selection with 1 μg/ml puromycin. The cells were treated with 1μg/ml doxycycline for at least 72 hours for the induction of knockdown. The RNA interference (RNAi) efficiency was assessed by Western blot analysis using RecQ1 antibody.

### IdU labeling and immunoprecipitation of replicated DNA

IdU labeling was performed as described previously (54). Briefly, cells were pulsed with 30 μM of IdU for 30 min, washed twice with cold PBS, and the episomal DNA was extracted by a modified Hirt’s method. The samples were dissolved in 500 µl TE (10 mM Tris-HCl, 1 mM EDTA), sonicated to get an average length of 700 bp, and heat denatured at 95 °C for 5 min. 10% of the extracted DNA was used as input control, and 1 μg of the mouse of anti-IdU antibody was added to samples and incubated at room temperature with constant rotation for 1 h. Magnetic Protein A/G Antibody was used to pull down the bound IdU labeled DNA, and the beads were washed once with 1X IP buffer (10 mm NaPO4 pH 7.0, 140 mM NaCl, and 0.05% Triton X-100), resuspended in 200 μl of lysis buffer (50 mM Tris-HCl (pH8.0), 10 mM EDTA, 0.5% SDS, 0.25 mg/ml Proteinase K) and incubated overnight at 37 °C for elution. This was followed by adding 100 μl of lysis buffer and incubating at 50 °C for 1 h. The eluted DNA was phenolized and precipitated for the quantitation of IdU-labeled DNA in a real-time PCR by amplifying the oriLyt region.

### KSHV Virion purification

KSHV virions were purified as described previously (57). Briefly, 15 million cells were induced with 0.3M NaB and 20 ng/mL TPA for 96 hours; culture supernatant was cleared by centrifugation and filtered through a 0.45-μm filter to remove cell debris. The virus was concentrated by ultracentrifugation (25,000 rpm for 2h at 4°C), resuspended in serum-free RPMI, and 50 μl of virus supernatant diluted with 250 μl of 1X PBS was digested with DNase-I (5 U/50 µl of supernatant) at 37°C for 1 h. DNase-I was heat inactivated at 70°C for 10 min, and supernatants were mixed with an equal volume of lysis buffer (0.1 mg/ml of proteinase K in water) and incubated at 50°C for 1 h. Proteinase K was heat-inactivated at 75°C for 20 min, and DNA was purified using PCI (Phenol: Chloroform: Isoamyl alcohol), precipitated with ethanol at -20°C and resuspended in sterile Milli-Q water. The viral DNA was amplified with qPCR using primers specific for LANA, and virus quantities were calculated based on the LANA standards. All the reactions were run in triplicates.

### Quantitation of KSHV RNA

Total RNA was isolated from the cells using Illustra RNAspin Mini Kit (GE Healthcare) according to the manufacturer’s protocol. cDNAs were generated using a high-capacity RNA-to-cDNA kit (Applied Biosystems Inc.) per the manufacturer’s protocol and quantified using specific primers. The Ct values were normalized to the housekeeping gene, GAPDH. All the reactions were run in triplicates.

### Determination of viral genome copies

Five million KSHV-positive cells were induced for lytic reactivation for 24 hours with or without compound treatment. Total genomic DNA was isolated using PureLink™ Genomic DNA Purification Kit according to the manufacturer’s instructions and quantified using primers specific to the DNA sequence of RTA. The Ct values were normalized to the housekeeping gene, GAPDH. All the reactions were run in triplicates.

### Statistical analysis

P-values were calculated by a two-tailed t-test using GraphPad Inc. (Prism 8) software for statistical significance. Asterisks represent the *P* value < 0.05 (*), *P* value < 0.01 (**) and <0.001 (***), where ns denotes non-significant

## RESULTS

### oriLyt DNA formed stable G4 structures

DNA G-quadruplexes are secondary structures formed in G-rich stretches of DNA that have been implicated in several biological processes. These structures have been reported in the latent nuclear protein of EBV, terminal repeat region of KSHV, viral gene promoters, and in the oriLyt region of the Human Cytomegalovirus (HCMV) (52, 58, 59). Given the critical role of the G4 structures in the viral life cycle, we were interested in investigating their role in the lytic DNA replication of KSHV. Firstly, we analyzed the oriLyt sequence of KSHV for the presence of G-quadruplex using a web-based tool, QGRS mapper, that predicts the formation of these structures (http://bioinformatics.ramapo.edu/QGRS/index.php) (60). G-scores, a predictor of the G-quadruplexes formation, were determined for the oriLyt region of KSHV, and the region with the high G-scores is represented by a yellow highlighted region (Fig. 1A). The oriLyt sequence with the highest G-score was located between the AT-rich region and the RRE element. Since G4 structures are unique concerning their structure and folding (Fig. 1B), they display distinct biophysical and biochemical properties. We validated the formation of G-quadruplexes by performing CD spectroscopy of an oligo containing the wild-type oriLyt G4 site or a scrambled oligo. Oligo with specific G4 site showed a spectral pattern with a maximum at 260 nm and a negative minimum at 240 nm for wild-type oriLyt oligo (Fig. 2A panel a). This pattern is specific for G-quadruplexes and was not observed for scrambled oligo (Fig. 2A panel b). We further confirmed the G-quadruplex formation on the oriLyt oligo in an electrophoretic mobility shift assay. The wild type and scrambled oriLyt DNA oligos were labeled with ^32^P using Terminal deoxynucleotidyl transferase and resolved in the presence of K+ ions on a 15% native gel, following which autoradiography was used to determine the mobility of these oligos (Fig. 2A panel a). Our results showed that due to the formation of condensed G-quadruplex structures, wild-type oriLyt G4 forming DNA oligo migrated faster than the scrambled oligo (Fig. 2A panel b). To corroborate that the increased mobility of the wild-type oriLyt oligo was caused by the formation of these secondary structures, we used antisense oligos to disrupt the formation of G-quadruplexes. The antisense oligos (AS1: GCCACCGAACAACCCC and AS2: CACTAGCCCCCCCC) were complimentary to the G-rich region of wild-type oriLyt oligo. Following the incubation of these antisense oligos with G4 oriLyt oligo and resolution on a native gel, we observed a shift in the mobility of wild-type oriLyt oligo in the presence of specific antisense oligos as compared to the wild-type oriLyt oligo alone (Fig. 2A panel b, compare lane 3 with lane 1). Based on these findings, we concluded that the fast migration of oriLyt G4 oligos on the native gel is due to the formation of G-quadruplexes as the disruption of these structures by antisense oligos led to a significant shift in the mobility of the wild-type oligo. These results validated that oriLyt forms G-quadruplexes.

**Figure 1.**
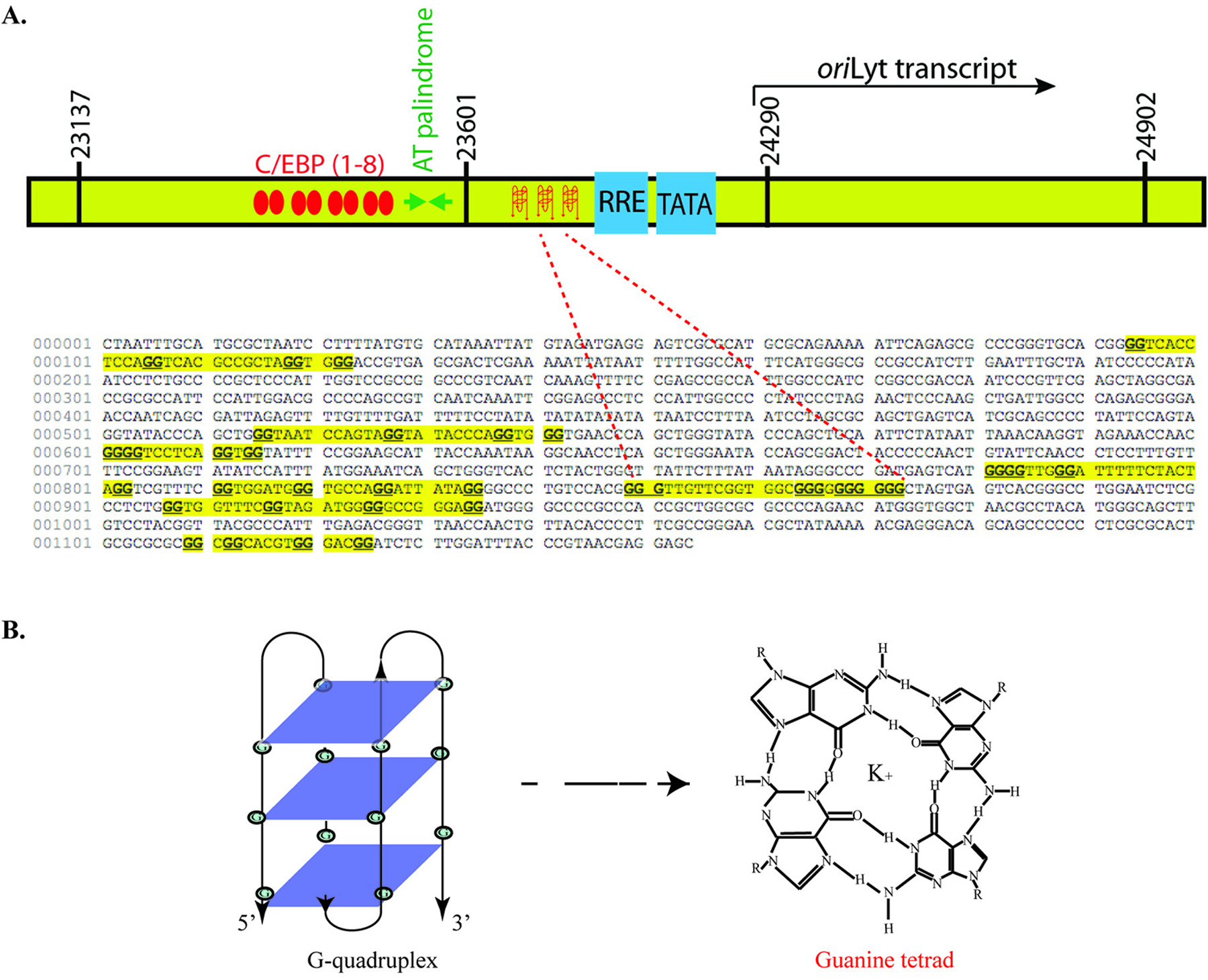
oriLyt region formed stable DNA G-quadruplexes. A. Schematic of the oriLyt region showing different domains, with the G-quadruplex (G4) forming regions highlighted in yellow. oriLyt sequence was imported into QGRS mapper software, which predicts G-quadruplex formation as a function of G-score. B. A guanine tetrad shows Hoogsteen hydrogen bonding between four guanine residues stabilized by a potassium ion in the center. Three guanine tetrads stack on top of each other to form a G-quadruplex.

**Figure 2.**
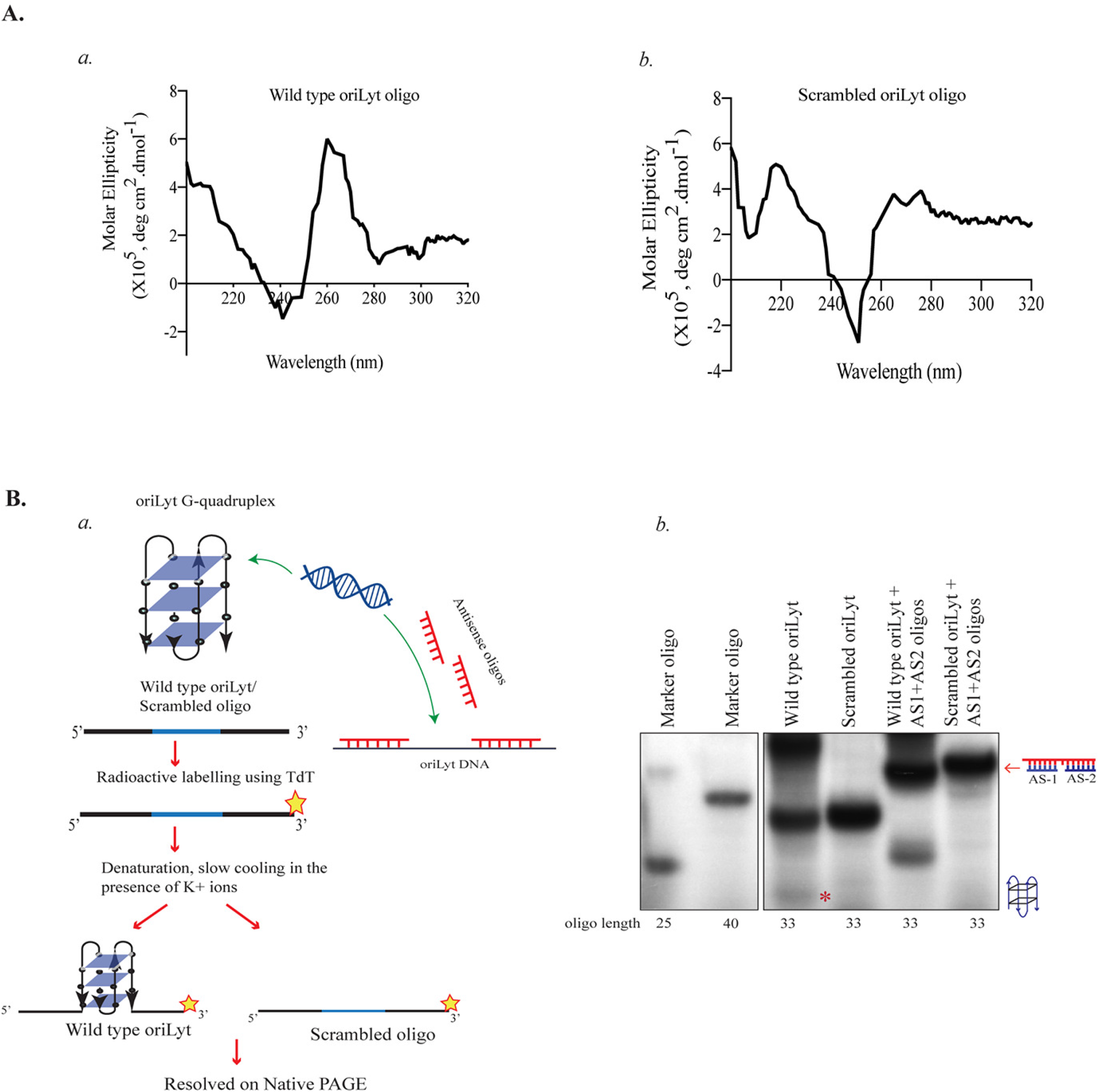
Biophysical and biochemical analysis confirmed the presence of G-quadruplexes in the oriLyt region. A. Circular Dichroism spectral analysis of *(a)* oriLyt wild type oligonucleotide having a high G-score and *(b)* Scrambled DNA oligonucleotide which could not form G4s was used as a control. The oligos were scanned for a wavelength range of 320 nm to 200 nm, and molar ellipticity was plotted on the Y-axis with a wavelength on the X-axis. B. *(a)* Schematic of steps involved in Native PAGE analysis of wild type oriLyt or scrambled oriLyt. *(b)* Electrophoretic mobility shift assay (EMSA) was performed in the presence of K+ ions on wild type oriLyt and scrambled oligos labeled with ^32^P and resolved on a native polyacrylamide gel. Antisense oligos (AS1 and AS2) complementary to wild-type oriLyt oligo, added in molar excess, were used in the indicated lanes to confirm that the G-quadruplex forming sequence caused the mobility shift.

### RecQ1 bound to G4 forming site on oriLyt

In light of the role of G-quadruplexes in cellular metabolism, it can be assumed that G4 structures interact with a variety of cellular proteins that either supports the formation or destabilize these structures, which is an area of extensive study (17). Interestingly, a study identified the involvement of RecQ1, a cellular helicase, in supporting KSHV lytic DNA replication by binding to the oriLyt region, an interaction facilitated by other viral proteins. RecQ proteins have been shown to facilitate DNA replication through their G4 resolving ability (61). This led us to investigate the G4 binding ability of RecQ1 with respect to oriLyt DNA. To achieve this, we first performed a DNA affinity pull-down assay, where wild-type oriLyt oligo carrying a G4 site and a scrambled oligo were biotinylated at the 3’ end using Biotin-11-UTP and Terminal deoxynucleotidyl transferase (TdT) followed by purification of biotinylated DNA and incubation with a cellular extract from KSHV positive BCBL-1 induced for 36 hours. After incubating the biotinylated DNA and the cell lysates together for three hours, streptavidin beads were added to capture the proteins associated with the DNA (Fig. 3A panel a). Beads were washed stringently to eliminate the non-specifically bound proteins, after which the proteins were resolved on SDS-PAGE.

**Figure 3.**
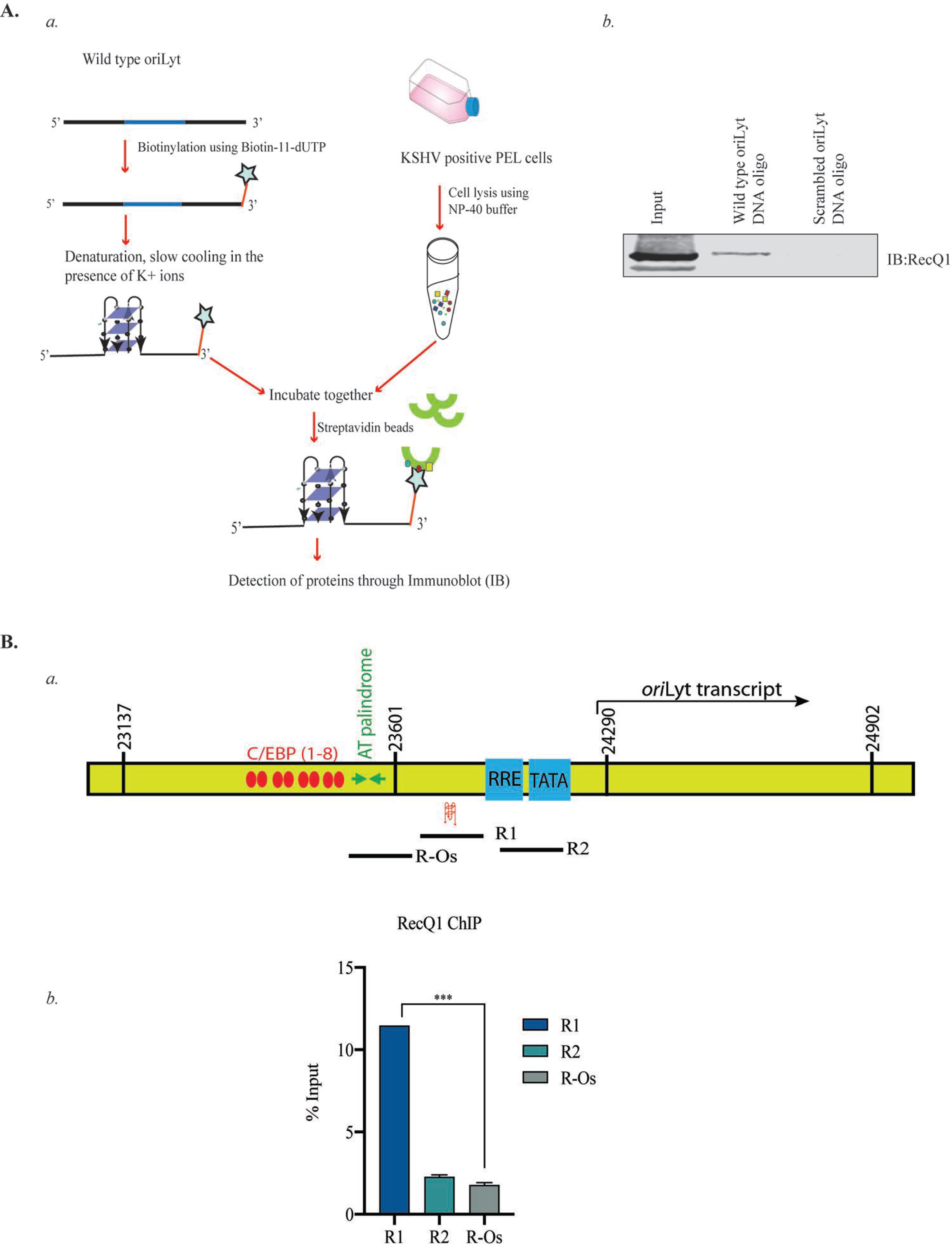
RecQ1 bound to the G-quadruplex forming site of the oriLyt. A. *(a)* Schematic of the steps involved in DNA affinity pull-down assay. *(b)* Immunoblot showing RecQ1 binding to the wild type oriLyt. Wild-type oriLyt and scrambled oligo (negative control) were biotinylated and incubated with cell lysate from lytically induced KSHV positive BCBL-1 cells, and streptavidin beads were added to pull down DNA-bound proteins followed by resolution of the samples on SDS-PAGE gel and detection by RecQ1 antibody. B. *(a)* Schematic of the oriLyt region depicting different regions (R1, R2, and R-Os) used for PCR amplification. *(b)* BCBL-1 cells were induced for lytic reactivation for 36 hours, following which the cells were harvested, crosslinked, and utilized to perform ChIP assay with RecQ1 antibody to determine the relative binding of RecQ1 at different regions (R1, R2, and R-Os) of oriLyt using specific primers.

Upon detection using the RecQ1 antibody, we observed that RecQ1 specifically pulled down with wild-type oriLyt oligo and not the scrambled oligo (Fig. 3A panel b). Next, we wanted to validate our findings *in vivo*, so we performed Chromatin immunoprecipitation assay using RecQ1 antibody on BCBL-1 cells that had been induced for lytic reactivation for 36 hours. Three sets of primers were used to quantify relative RecQ1 enrichment of ChIP DNA; R1 specific for the region that carries a G4 site, R2 specific for the region that was previously identified to bind oriLyt but does not possess a G4 site, and R-Os, specific for a region outside the RecQ binding area of oriLyt (50) (Fig. 3B panel a). Upon comparison, we observed that RecQ1 is selectively bound to R1 more than R2 and R-Os. Collectively, these findings revealed the binding of RecQ1 to the G4 site in oriLyt of KSHV (Fig. 3B panel b).

### RTA recruited RecQ1 to the oriLyt region

RTA plays an indispensable role in lytic DNA replication and interacts with another viral protein, K8, to perform various functions in replication initiation. RTA has been shown to bind to RRE in the oriLyt region, activate the promoter of oriLyt, and initiate transcription. In addition, it has been shown to interact with proteins of the prereplication complex and recruits them to the origin to facilitate lytic DNA replication (14). There has been evidence that RecQ1 binds to RTA directly, and its binding to the oriLyt region is compromised in the absence of the RRE domain. We wanted to investigate the binding of RTA and RecQ1 with respect to G4 forming sites in the oriLyt. To achieve this, we first performed a chromatin immunoprecipitation assay with RTA in 24-hour induced BCBL-1 cells and quantified RTA enrichments using the three primers mentioned above. Not surprisingly, we found that RTA was relatively enriched at R2, which contains the RRE domain but no G4 site (Fig.4.A panel a). Next, we analyzed the relative enrichment of RecQ1 at three regions in the absence of RTA. To this end, we performed a chromatin immunoprecipitation assay with RecQ1 in iSLKTet-RTABAC16-WT and iSLKTet-RTA-Bac16-RTASTOP cells. We observed that the relative binding of RecQ1 at R1 decreased in cells where RTA had been depleted compared to wild-type cells (Fig.4.A panel b).

**Figure 4.**
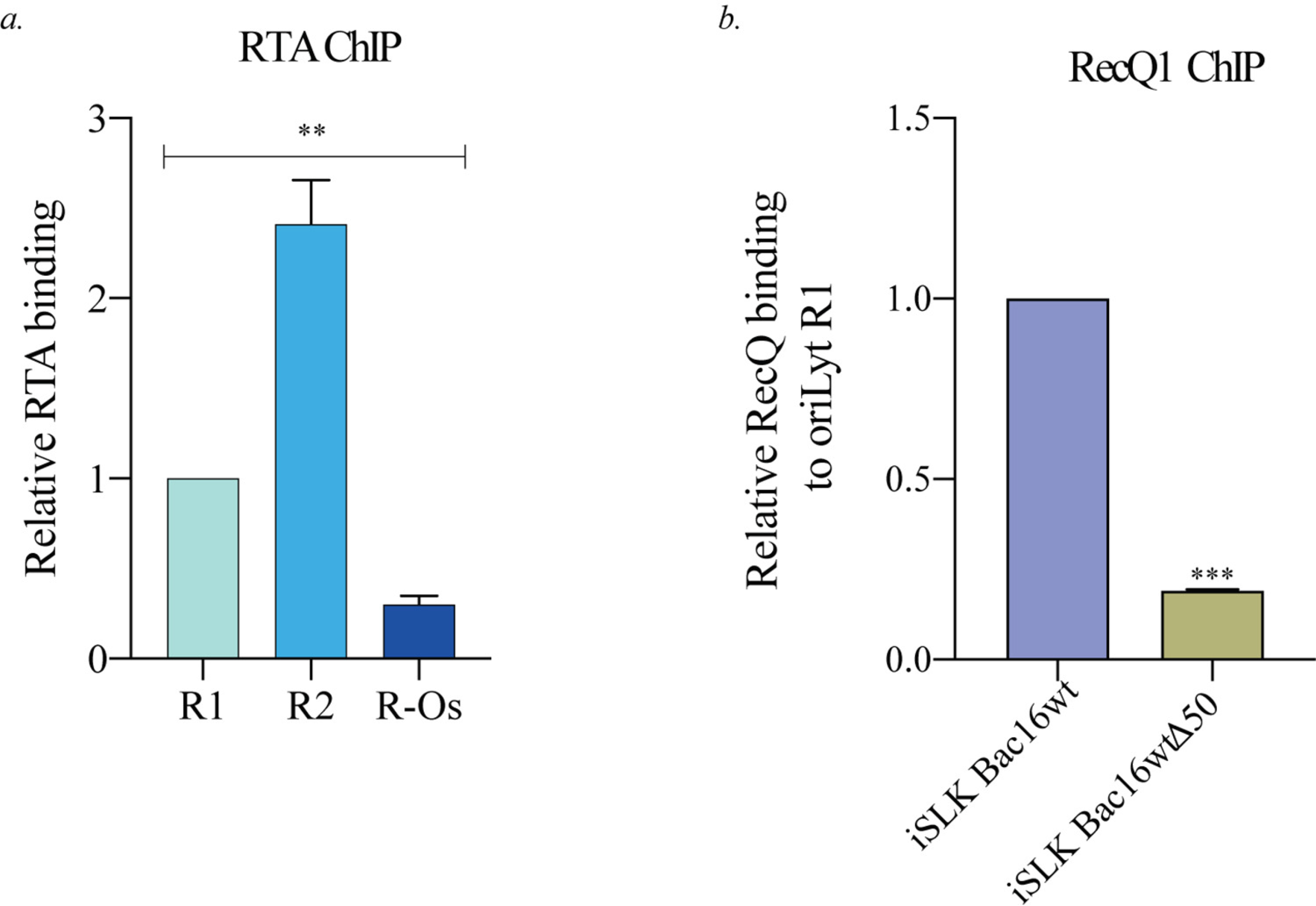
RTA facilitated the binding of RecQ1 at oriLyt. A. *(a)* ChIP assay was performed using lytically induced BCBL-1 cells, where the cells were harvested, crosslinked, and chromatin-bound DNA was pulled down using RTA antibody to determine the relative enrichment of RTA at different regions of oriLyt (R1, R2 and R-Os) using specific primers. *(b)* ChIP assay was performed using lytically induced iSLK-Tet-RTABAC16-WT and iSLKTet-RTA-Bac16-RTASTOP cells where the cells were harvested, crosslinked, and chromatin-bound DNA was pulled down using RecQ1 antibody to determine the relative binding of RecQ1 on oriLyt using primers specific for R1.

### Inhibition of RecQ1 helicase activity and RecQ1 depletion inhibits lytic DNA replication

Given the presence of RecQ1 at oriLyt and in the viral replication compartment from our observation and a previous report, we wanted to study its role in lytic replication intricately. We treated KSHV-positive cells with *N*-methyl meso-porphyrin IX (NMM), an inhibitor of RecQ1 helicase activity, and tested its effect on lytic replication.

First, we analyzed the amount of newly replicated DNA in compound-treated cells through IdU immunoprecipitation assay, where KSHV-positive cells were treated with NMM or DMSO and induced for lytic reactivation for 24 hours, following which the cells were labeled with IdU. Nascent DNA was pulled down using an anti-IdU antibody. The amount of newly replicated DNA was quantified using oriLyt-specific primers. Upon comparison, we observed that the amount of newly replicated DNA was reduced in cells treated with NMM compared to control cells treated with DMSO (Fig.5.A panel a). Since RecQ1 inhibition was negatively affecting active lytic replication, we then wanted to analyze the effect of RecQ1 inhibition on viral lytic gene expression and viral persistence. This was tested by quantifying transcripts of lytic immediate-early and late genes, i.e., ORF59 and ORF 8, respectively. NMM-treated cells showed a reduction in late gene transcription with no significant effect on early gene and latent gene transcription (Fig.5.A compare panels b, c, and d). Upon observing the negative impact of NMM treatment on late gene transcription, we wanted to analyze its effect on viral persistence. For this, we extracted genomic DNA and quantified the relative genome copies of the virus in DMSO or NMM-treated induced BCBL-1 cells, which showed a reduction of genome copies in cells treated with NMM compared to control cells (Fig.5.A panel e). Having established that RecQ1 activity inhibition led to the inhibition of lytic DNA replication and viral persistence, we wanted to test its effect on virion production. As expected, we observed a decrease in the virus titer in cells treated with NMM compared to control cells (Fig.5.A panel f).

**Figure 5.**
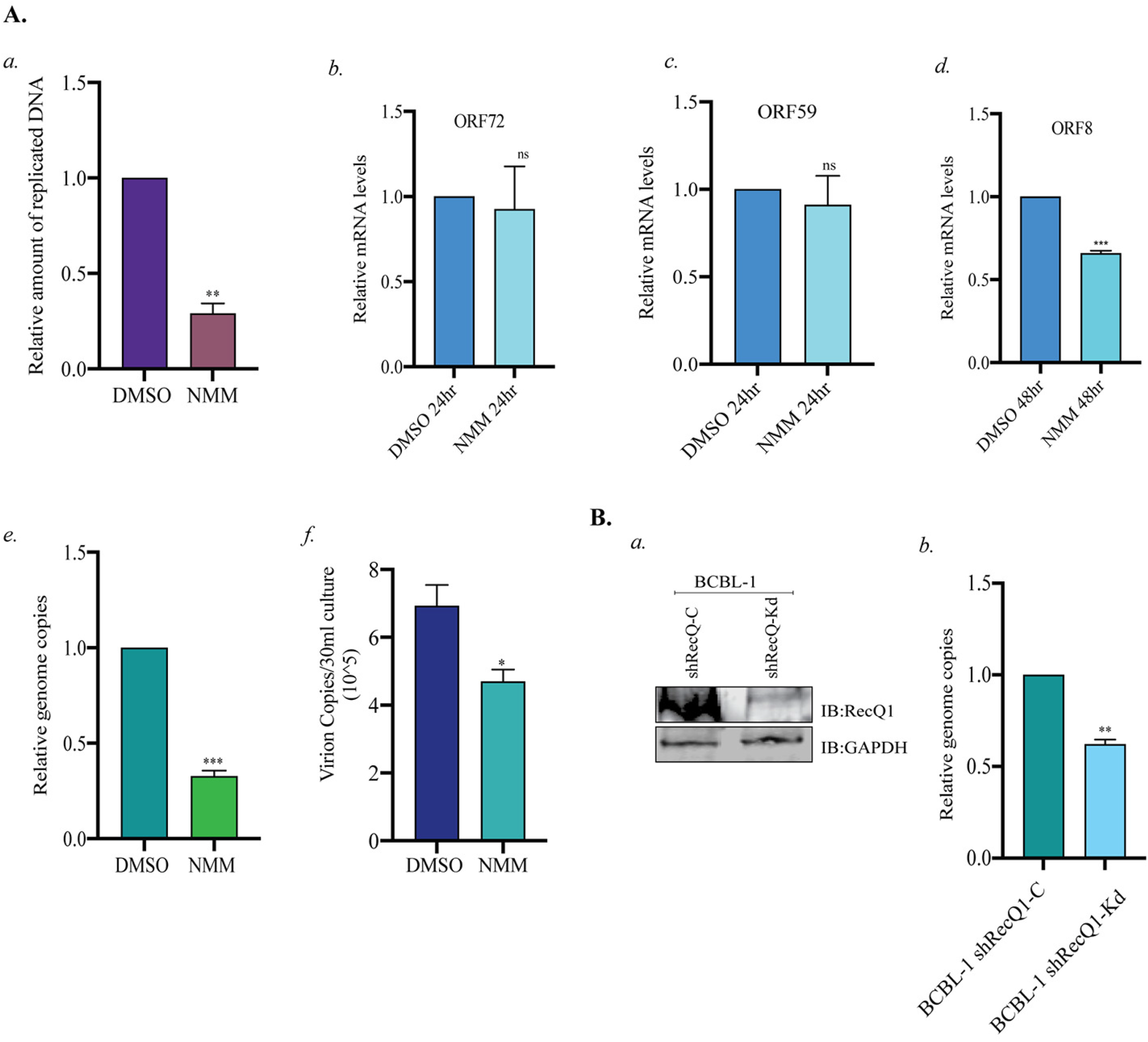
RecQ1 is essential for lytic DNA replication. A. BCBL-1 cells were treated with DMSO or 2uM NMM and induced for lytic reactivation *(a)* 24 hours post lytic induction; cells were labeled with IdU, genomic DNA was extracted, newly replicated DNA was pulled down anti-IdU antibody, and the DNA was quantified using primers specific for oriLyt region. *(b)* 24 hours post lytic induction, mRNA was extracted, cDNA was synthesized and quantified using latent gene, ORF72 specific primers *(c)* 24 hours post lytic induction, mRNA was extracted, cDNA was synthesized and quantified using immediate early lytic gene, ORF59 specific primers. *(d)* 48 hours post lytic induction, mRNA was extracted, and cDNA was synthesized and quantified using late lytic gene, ORF8 specific primers. *(e)* 24 hours post lytic induction, genomic DNA was extracted, and viral genome copies were quantified using primers specific for genomic DNA sequence of RTA. *(f)* 96 hours post lytic induction; the virus was concentrated from the supernatant of induced cells and quantified using ORF73 primers. B. *(a)* Immunoblot showing depletion of RecQ1 in control (shRecQ1-C BCBL-1) and knockdown (shRecQ1-Kd BCBL-1) cells 24 hours post lytic induction. *(b)* shRecQ1-C BCBL-1 and shRecQ1-Kd BCBL-1 cells were induced for lytic reactivation for 24 hours, following which genomic DNA was extracted, and viral genome copies were quantified using primers specific for the genomic DNA sequence of RTA.

All these findings led to the conclusion that RecQ1 unwinding activity was essential for KSHV lytic DNA replication as its inhibition negatively affected the amounts of actively replicated DNA, genome persistence, and virus titer. Similarly, we also tested the relevance of RecQ1 to lytic DNA replication through shRNA-based depletion of RecQ1. BCBL-1 cells stably transduced with shRecQ1 lentivirus were assayed for the depletion of RecQ1, post doxycycline treatment, where they showed a significant reduction in RecQ1 levels (shRecQ1-Kd) as compared to control cells (shRecQ1-C) (Fig.5.B panel a). Quantifying genome copies in shRecQ1-C and shRecQ1-Kd BCBL-1 cells showed reduced viral genome persistence in cells where RecQ1 was depleted (Fig.5.B panel b).

### G4 disruption inhibited RecQ1 binding and lytic replication

The presence of G4 sites in oriLyt and the binding of RecQ1 to these sites prompted us to evaluate the functional relevance of these sites in the genome through their disruption. The G-quadruplex formation is highly dependent on the occurrence of four repeats of at least three G-residues separated by a few bases in between. We performed site-directed mutagenesis in the oriLyt plasmid (8088sc), which changed the G-rich sequence from ACGGGGTTGTTCGGTGGC<u>GGGGG</u>GGGGGGGG (8088wt) to ACGGGGTTGTTCGGTGGC<u>AATAA</u>GGGGGGGG (8088mut), which reduced the propensity of formation of G4 structures indicated by a drop in G-score. (Red fonts show G4 sites) We tested these clones for their efficiency in lytic DNA replication by performing a transient replication assay using these plasmids. A transient replication assay was performed by transfecting BCBL-1 cells with 8088wt or 8088mut plasmids, and 24 hours post-transfection, the cells were induced for lytic replication for 48 hours. Genomic DNA was extracted from these cells, followed by digestion with EcoRI to linearize or DpnI and EcoRI to determine the replicated copies after Southern hybridization. Upon quantifying DpnI resistant/replicated DNA band, we observed lower lytic replication in cells transfected with 8088mut plasmid than in cells transfected with 8088wt plasmid (Fig.6.B panel a).

**Figure 6.**
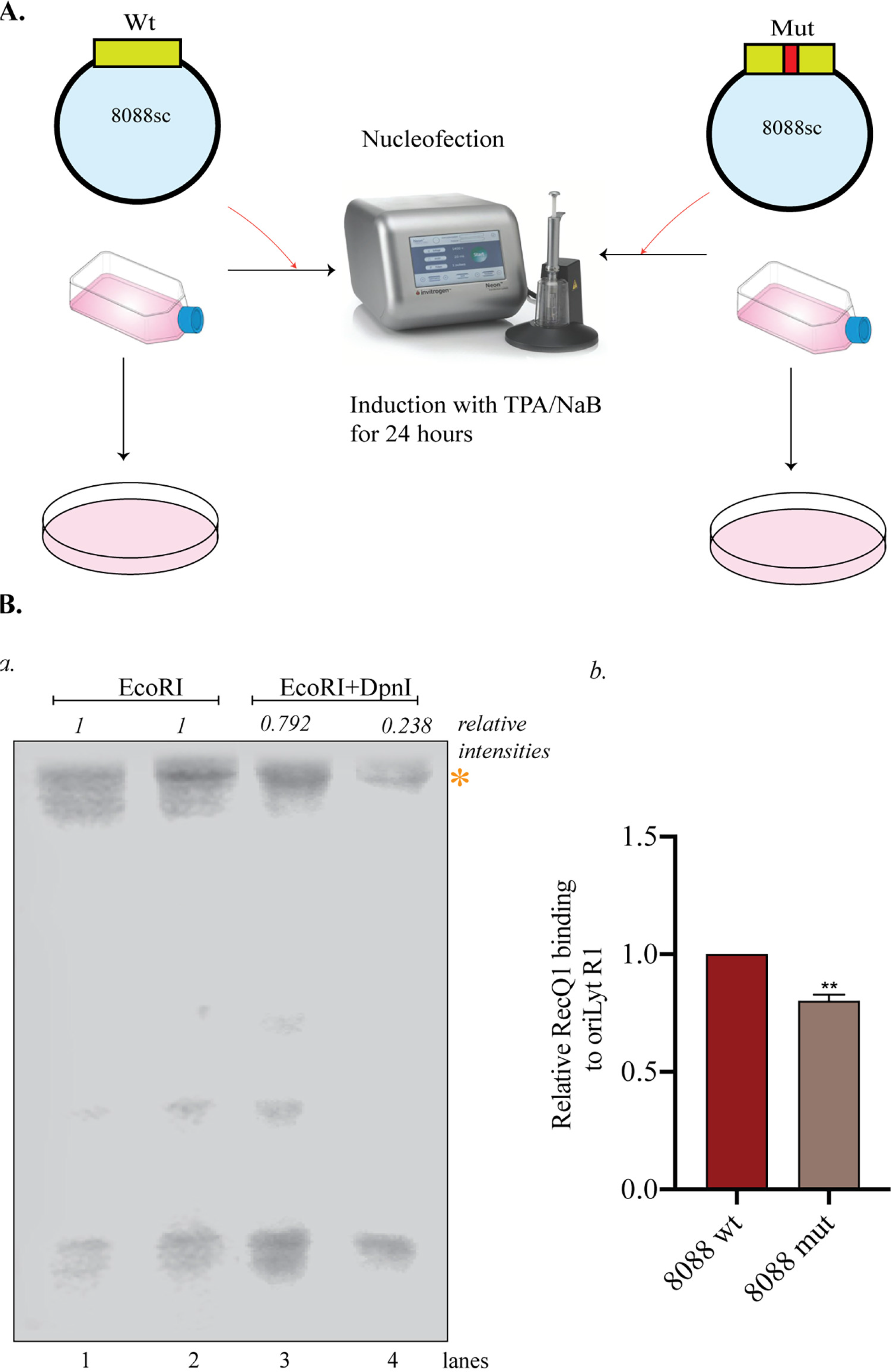
Disruption of G-quadruplex formation inhibited lytic replication and reduced RecQ1 binding to the G4 sites. A. *(a)* Schematic showing transfection of BCBL-1 cells with 8088wt or 8088mut plasmid. B. *(a)* BCBL-1 cells were transfected with 8088wt or 8088mut plasmid and induced for lytic reactivation. 48 hours post-induction, total genomic was extracted using Hirt’s procedure and digested with either EcoRI or EcoRI and DpnI. DNA was subjected to Southern blotting and probing with an 8088sc specific probe. *(b)* ChIP assay was performed using 8088 wt or mut transfected and 24-hour lytically induced BCBL-1 cells, where the cells were harvested, crosslinked, and chromatin-bound DNA was pulled down using RecQ1 antibody to determine the relative enrichment of RecQ1 at R1 region of oriLyt using specific primers.

In our previous experiments, we showed that RecQ1 binds explicitly to the oriLyt G4 site compared to other regions in oriLyt. Since mutation of G4 sites deterred the formation of G4 structures (indicated by the G-scores), we wanted to test the effect of disruption of G4 on RecQ1 binding. We achieved this by performing a Chromatin Immunoprecipitation assay on BCBL-1 cells transfected with 8088wt or 8088mut using RecQ1 antibody. BCBL-1 cells were transfected with 8088wt or 8088mut plasmids, and 24 hours post-transfection, the cells were induced for lytic reactivation for 24 hours. ChIP was performed using the RecQ1 antibody, and the immunoprecipitated DNA was amplified using primers specific for R1. The relative binding of RecQ1 to R1 was reduced in cells transfected with 8088mut compared to cells transfected with 8088wt, indicating that RecQ1 selectively binds to the G4 sites, and this binding is compromised by mutation of the G4 sites (Fig.5.B panel b).

### G-quadruplex stabilization reduced the initiation of lytic replication

Even though many G4 sites have been predicted in the genomes, not all will result in stable G-quadruplex formation *in vivo*. Many G-quadruplex ligands have been discovered that bind G4 sites selectively and stabilize them. G4 sites have been associated with replication initiation in previous reports and a recent study that established a link between G4 sites and origin activity in the mammalian genome. This has prompted us to study the effect of the formation of these G4 structures on lytic DNA replication initiation in KSHV(61, 62).

Herpesviruses have been proposed to replicate via bidirectional theta-type replication and rolling circle replication, ultimately resulting in the generation of viral concatamers. However, the exact mechanism is not yet known (63). We analyzed the effect of G-quadruplex stabilizing ligands on replication initiation through Single Molecule Analysis of the Replicated DNA (SMARD) assay, a technique used to study replication events in EBV and KSHV (64, 65). This involves multiple steps beginning with consequent labeling of cells with halogenated nucleotides, IdU and CldU, embedding the cells in agarose plugs, lysis of cells, and digestion and linearization of DNA. The plugs containing DNA are resolved on Pulse field Gel electrophoresis, where a band specific for the KSHV genome is excised, and DNA is extracted from the agarose plugs following gelase digestion. This is followed by the stretching of DNA on positively charged glass slides and fluorescent *in-situ* hybridization (FISH) with biotinylated probes. The labeled DNA is finally detected by monoclonal antibodies against IdU and CldU and fluorescently labeled secondary antibodies, whereas the biotinylated probes are detected by fluorescence-conjugated avidin. The DNA molecules used to identify replication initiation sites possess labels from the nucleotide analogs (IdU: red and CldU: green) and were arranged based on the increasing length of the IdU label. The molecules signifying a bidirectional initiation site will show a progressively increasing red signal surrounded by the green signal, indicating the replication fork’s bidirectional movement. Upon analysis of labeled DNA molecules from DMSO or TMPyP4 treated BCBL-1 cells, we observed a reduction in initiation of DNA replication denoted by the absence of DNA molecules specific for origin, which are characterized by red signal flanked by green signal (Fig.7.). Thus, it can be concluded that G-quadruplex stabilization leads to a reduction in lytic DNA replication initiation.

**Figure 7.**
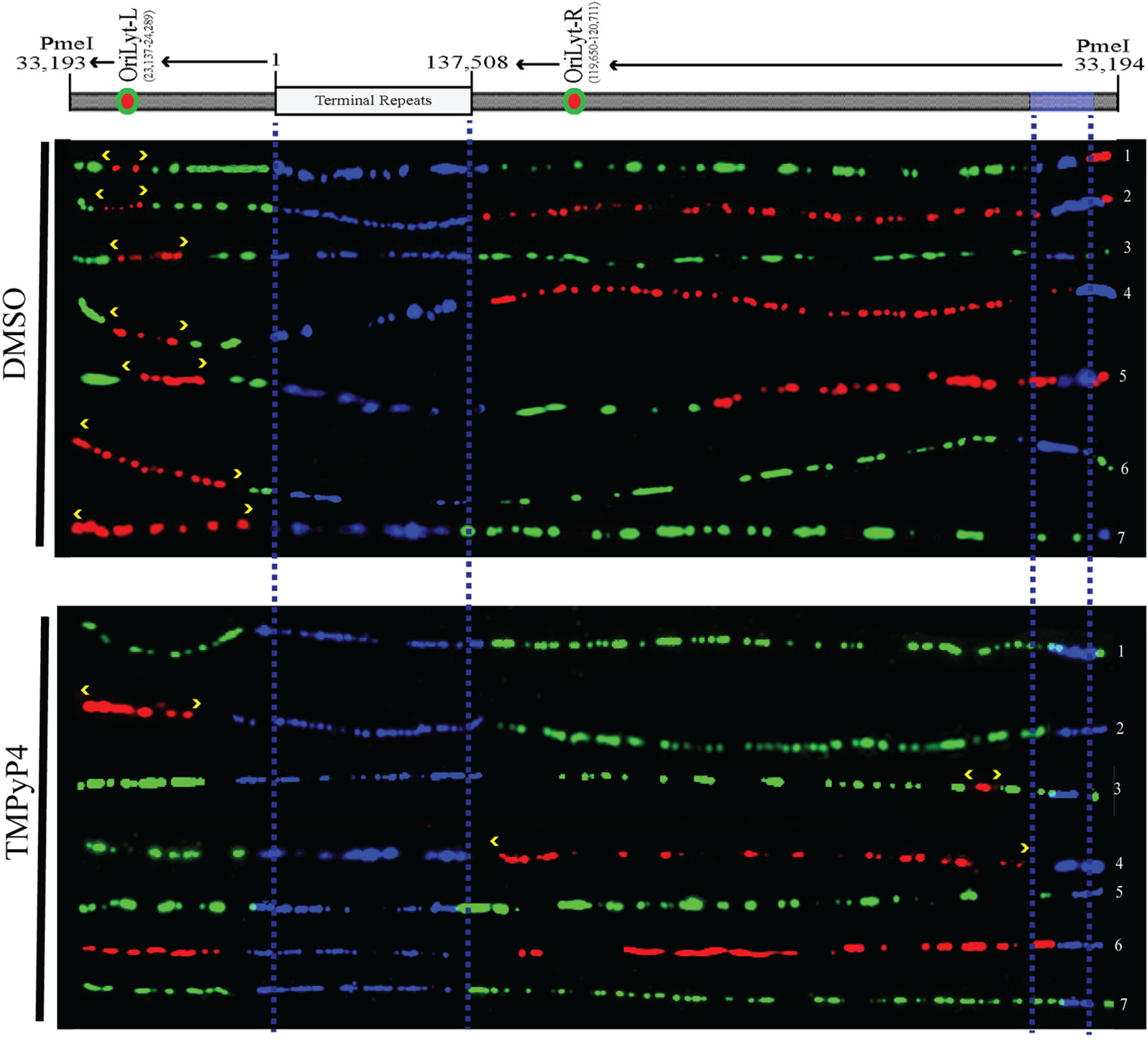
G-quadruplex stabilizing agent reduced initiation of lytic DNA replication from oriLyt. BCBL-1 cells were treated with DMSO or TMPyP4, induced for lytic reactivation for 24 hours, and SMARD was performed on them. KSHV genome was detected by blue FISH probe signals. Red and green labels represented IdU (first label) and CldU (second label), respectively, following detection by immunostaining. Yellow arrows denoted actively replicating DNA from the origin, i.e., oriLyt L, and green labels flanking red showed the bidirectional movement of the replication fork. The progressions of replication forks are marked with yellow arrows.

### G-quadruplex stabilization inhibited lytic DNA replication

Through our previous experiments, we have shown that G4 structures regulate lytic DNA replication. To evaluate the role of G4 formation in virus lytic replication, we used two G-quadruplex stabilizing compounds, PhenDC-3 and TMPyP4, to treat BCBL-1 cells and DMSO as a negative and PAA, a known inhibitor of lytic replication as a positive control.

G4 stabilization inhibited the initiation of lytic DNA replication, which is essential for the transcription of late lytic genes. Therefore, we wanted to analyze the effect of G-quadruplex formation on lytic gene transcription. This was tested by quantifying transcripts of lytic immediate-early and late genes, i.e., ORF59 and ORF65, respectively. Late gene expression was significantly reduced after treatment of cells with G4 stabilizing compounds (PhenDC-3 and TMPyP4) and PAA compared to DMSO control with no significant effect on early gene and latent gene transcription (Fig. 8.A compare panels a, b, and c). Next, we tested the effect of G4 stabilization on the persistence of the virus in the host cell by quantifying the relative viral genome copies in BCBL-1 cells treated with DMSO or other compounds. Following induction of the compound treated cells for lytic reactivation for 24 hours, genomic DNA was isolated, and viral genome copies were quantified using gene-specific primers. Strikingly, we observed a reduction in viral genome copies in BCBL-1 cells treated with Phen DC-3 and TMPyP4 along with cells treated with the positive control PAA in comparison with control DMSO-treated cells, leading us to the conclusion that stable G-quadruplex formation in the cells led to defective genome persistence (Fig. 8. A panel d). The next step was to assess the effect of reduction in genome copies and late gene expression on virion production. Compared to DMSO-treated cells, BCBL-1 cells treated with PhenDC-3 or TMPyP4, or PAA, demonstrated a significant reduction in the virus titer, which was consistent with our previous findings (Fig. 8.A panel e). All these observations suggested that the formation of G4 structures in the KSHV genome inhibits lytic replication of the virus.

**Figure 8.**
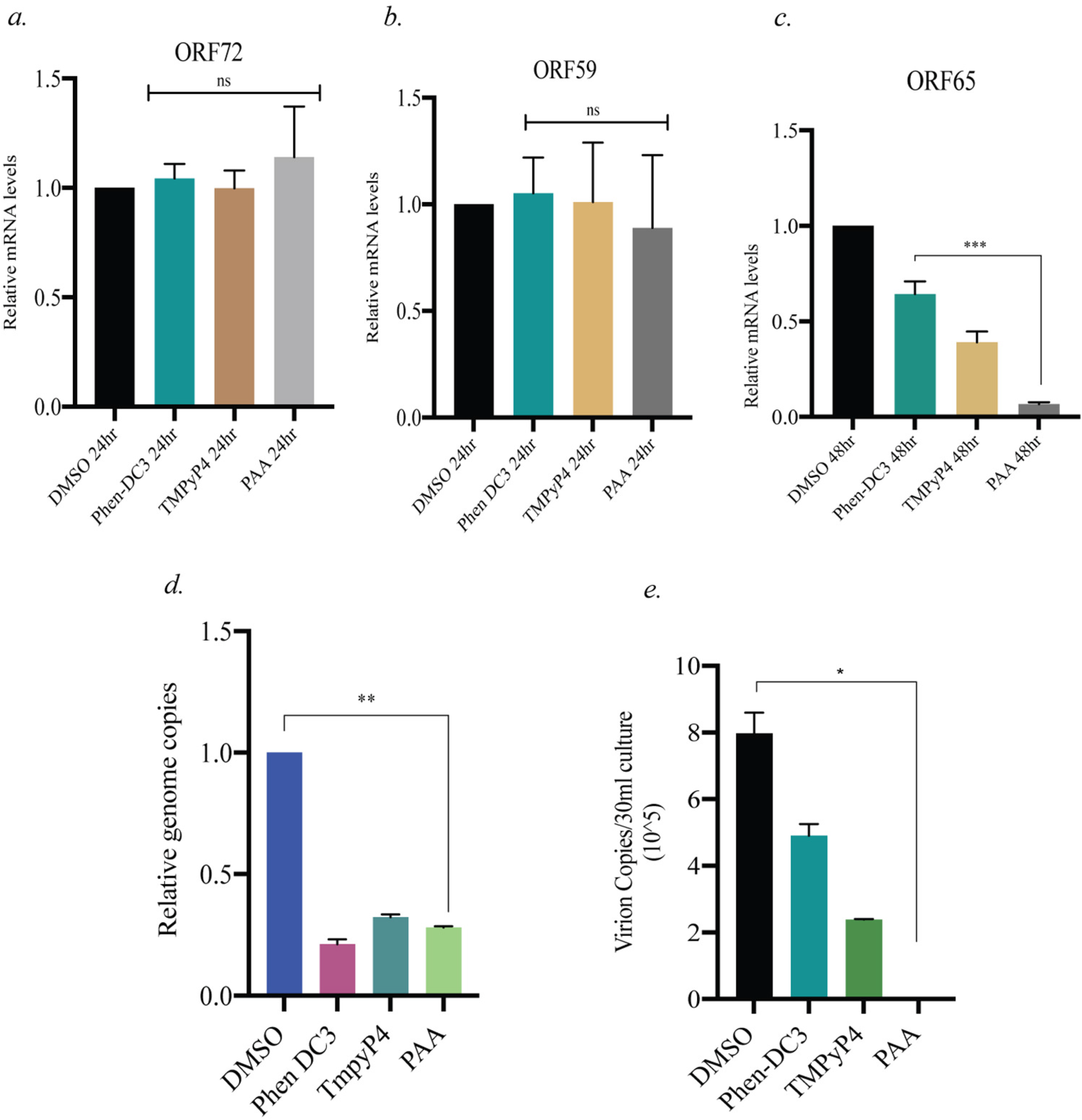
G-quadruplex stabilization inhibited lytic DNA replication. C. BCBL-1 cells were treated with DMSO/10μM Phen-DC3/10μM TMPyP4/0.5mM PAA and induced lytic reactivation. *(a)* 24 hours post lytic induction, mRNA was extracted, cDNA was synthesized and quantified using latent gene, ORF72 specific primers *(b)* 24 hours post lytic induction, mRNA was extracted, cDNA was synthesized and quantified using immediate early lytic gene, ORF59 specific primers *(c)* 48 hours post lytic induction, mRNA was extracted, cDNA was synthesized and quantified using late lytic gene, ORF65 specific primers. *(d)* 24 hours post lytic induction, genomic DNA was extracted, and viral genome copies were quantified using primers specific for genomic DNA sequence of RTA. *(e)* 96 hours post lytic induction; the virus was concentrated from the supernatant of induced cells and quantified using ORF73 primers.

**Figure 9.**
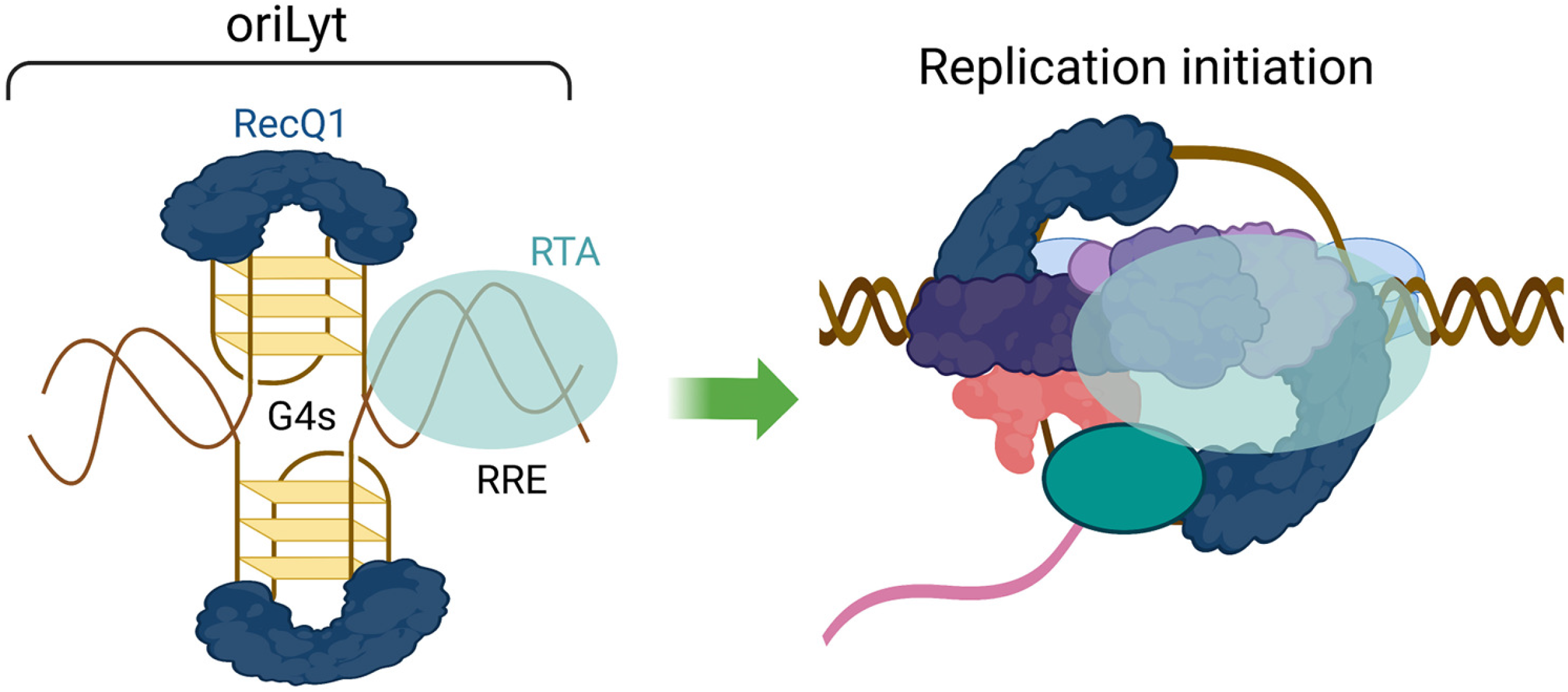
Potential model of G-quadruplex mediated replication initiation at KSHV oriLyt. The RecQ1 (DNA helicase) binds to the G-quadruplex sites of the oriLyt to potentially unwind the DNA structure and facilitate DNA replication, assisted by the binding of other proteins, including RTA.

## Discussion

KSHV, a member of the gamma herpesvirus family, maintains a biphasic life cycle consisting of latent and lytic cycles. Whereas latency ensures the lifetime persistence of the virus in the host, lytic DNA replication is essential to maintain viral reservoirs in infected cells, disseminate the virus and promote tumorigenesis through the expression of oncogenic gene products (6).

Latent DNA replication occurs synchronously with the cell cycle, and the duplicated genome is segregated into daughter cells. LANA is responsible for these functions in addition to latent DNA replication, where it recruits cellular DNA replication proteins to the origin of latent replication, oriP (66). Latency to lytic reactivation switch can be triggered by various stimuli such as Hypoxia, oxidative stress, viral coinfection, and chemicals (67). RTA is indispensable for lytic reactivation and is responsible for the switch of the viral life cycle from latency to lytic cycle through activation of its promoter and promoters of other viral genes, facilitates viral DNA replication through binding to the origin, acting as a component of the prereplication complex and recruitment of cellular and viral factor to the origin, all of which are required for lytic DNA replication (14). This mode of replication differs from the latent mode in many ways as it originates from a distinct origin of replication, oriLyt, which requires the functioning of multiple viral proteins and results in the production of thousands of viral genomes. Apart from viral proteins, several cellular proteins have been shown to aid lytic DNA replication and bind to the oriLyt region, which includes topoisomerases (Topo) I and II, RecQ1, poly (ADP-ribose) polymerase I (PARP-1), DNA-PK and scaffold attachment factor A (SAF-A) to name a few (50). G-quadruplexes are regulatory structures formed in G-rich DNA/RNA sequences that have been reported to play significant roles in biological processes. These structures have been identified in viral genomes such as HIV, EBV, HPV, HCMV, and KSHV (59, 68). Owing to the high GC content in their genomes, Herpesviruses genomes could be expected to have a high propensity for forming G4 structures. Many studies have reported high putative quadruplex sequences (PQS) in Herpesvirus genomes (18, 59, 69). The regulation of DNA replication by G4 structures has been well studied, but their role in origin firing or replication initiation is relatively unexplored (61, 70, 71). A recent study provides strong evidence regarding the substantial role of G4s in the origin activity (62).

RecQ1, a cellular helicase, is a member of the RecQ helicase family, known for their roles in recombination, repair, and replication. In addition, other members of the RecQ family, such as BLM and WRN helicase, have been shown to unwind G4 structures and facilitate telomeric replication (61). RecQ1 has been shown to promote genome stability through its role in DNA damage response and facilitate restarting paused replication forks (72). Additionally, it has been proposed to have a role in origin activity and has been shown to participate in lytic DNA replication by binding to the oriLyt region of KSHV and EBV (50, 51, 71). We wanted to link the association of RecQ1 and oriLyt to the presence of G4s in the latter and perform studies to explore the relevance of this association in the KSHV life cycle.

As the TR region of KSHV has been demonstrated to form G4 structures, we explored the possibility of forming these structures in the oriLyt region. Software-based analysis of the DNA sequence of oriLyt reveals a strong possibility of the formation of G4 in a few regions, which was confirmed later through CD spectroscopy and EMSA. The involvement of RecQ1 in the prereplication complex and the viral replication centers through binding at oriLyt has shed light on the origin-based activity of RecQ1. G4 unwinding activity, another widely studied function of RecQ1, could explain its engagement at oriLyt. As an extension to previous studies about RecQ1 in KSHV, we found RecQ1 to be selectively enriched at the G4 forming site in oriLyt as compared to any other region, and this was confirmed biotinylated DNA pulldowns and through Chromatin Immunoprecipitation assay.

Moreover, the role of RTA in RecQ1-oriLyt binding was also substantiated by reduced pull-down of RecQ1 at oriLyt in RTA-deleted cell lines. A reduction in genome copies of the virus in RecQ1-depleted cells showed further evidence about the role of RecQ1 in lytic replication. More interestingly, the inhibition of helicase activity of RecQ1 through the use of NMM led to a reduction in actively replicated DNA, genome copies, virion concentration, and late gene expression. Additionally, we demonstrated the importance of multiple stretches of G-residues in G4 structures on lytic replication through mutation of these residues in oriLyt. We observed defective active lytic replication and reduced RecQ1 binding to the oriLyt region.

Importantly, we evaluated the effect of G4 stabilization on lytic replication by using compounds like PhenDC-3 and TMPyP4, which promote G4 formation. Our particular interest was the observation through SMARD, which demonstrated that the cells in which G4 formation had been stabilized displayed compromised initiation of replication from the origins. These results were also corroborated in the subsequent assays, where G4 stabilization led to a reduction in viral genome copies, late lytic gene expression, and progeny virion production. Taken together, our findings indicate that G-quadruplex formation in the oriLyt region regulates the initiation of lytic DNA replication in KSHV, and RecQ1 binding at the G4 sites promotes replication, possibly through unwinding these secondary structures. Still, more studies need to be performed to shed light on the mechanism associated with RecQ1-mediated oriLyt replication.

## Acknowledgments

We thank Dr. Jae Jung (University of Southern California) for iSLK-Bac16-TetRTA and iSLK-Bac16-ΔRTA cell lines. We also thank Dr. Sudha Sharma (Howard University) for the RecQ1 antibody. We also acknowledge Dr. Matthew Tucker (University of Nevada, Reno) for help with CD spectroscopy. This work was supported by the National Institute of Health, grant number AI105000.

